# Genome-wide associations of host susceptibility to helminth and blood pathogens in spatially structured rodent populations

**DOI:** 10.64898/2026.05.19.726205

**Authors:** Abosede E. Olarewaju, Jarosław Bryk, Victoria I. Ayansola, Alyssa Dunn, Anna Rybińska, Agnieszka Kloch

## Abstract

Parasites are ubiquitous drivers of host evolution by exerting strong selective pressure on natural populations. Understanding the genetic basis of host susceptibility to infection is important to know how host-pathogen interactions shape patterns of resistance and diversity in natural populations. We conducted a genome-wide association study (GWAS) to identify host genetic variants associated with infection by helminth and blood pathogens in spatially structured populations of Bank voles (*Myodes glareolus;* (Schreber, 1780). We genotyped 182 individuals sampled from ten sites in central Europe using quaddRAD sequencing, retaining 30,206 high-quality single-nucleotide polymorphisms (SNPs). Associations between SNP genotypes and parasite infection status were tested using mixed models controlling for relatedness, with host body mass included as a covariate. Across parasite taxa, we identified twelve SNPs exceeding genome-wide significance with the strongest signals detected for the intestinal nematode *Heligmosomum mixtum*. The variants identified are all intergenic, intronic, upstream or downstream of genes, with none predicted to alter coding sequences. These genes are not classical immunity genes but some are implicated in cytokine production, PI3K/AKT signalling and p38 MAPK pathway, suggesting that selective pressure from pathogens does not only act on known immunity genes, but on broader regulatory and metabolic networks. This finding suggests that variation in gene expression may be important for the differences in host susceptibility or resistance to parasitic infections.

## Introduction

Parasites are ubiquitous drivers of host evolution by exerting strong selective pressure on natural populations, which influences host survival and reproductive success (Duffy et al. 2008; Ebert & Fields, 2020). Wild populations are exposed to a wide range of parasites and pathogens that may trigger different immune responses in their hosts due to genetic variation at immune-related loci (Cheynel et al. 2023). Coevolution between hosts and parasites bases upon reciprocal genetic change, which manifests as an evolutionary arms race or Red Queen dynamics, expressed as antagonistic adaptations or continuous cycles of reciprocal adaptation, described in details in many studies (e.g. Filipiak et al. 2016; Papkou et al. 2016; Vorburger & Perlman, 2018).

In the wild, parasite communities are rarely homogeneous. Variation in parasite prevalence, genetic diversity, and community composition among host populations can generate spatially heterogeneous selection pressures, potentially promoting genetic differentiation and local adaptation in hosts (Minias et al. 2023; Phillips et al. 2021). Such parasite-mediated selection may contribute to host population divergence by favouring different alleles across environments or locations (e.g. Bonneaud et al. 2006, Kloch et al. 2010), although similar patterns of differentiation may also arise from neutral processes such as genetic drift and restricted gene flow. Disentangling adaptive genetic responses to parasites from neutral demographic structure therefore remains a major challenge in evolutionary ecology. The confounding effect of population structure is the key challenge in identifying genetic variants underlying susceptibility to infection. Differences in allele frequencies between populations could arise due to historic demographic processes and environmental factors, including parasite-driven selection pressure. Both the host’s genetic history and its exposure to parasites contribute to variation in susceptibility to infection in the host (Oyesola et al. 2022). For example, spatial differences in parasite communities can coincide with genetic differentiation among populations, making the interpretation of genetic associations difficult. Long-term ecological studies, such as those on bank voles, show that some populations may lack certain parasite species entirely, while others carry high parasite loads of the same parasite species (Grzybek et al. 2015). Not properly accounting for the effects of population structure could lead to spurious associations and reduce the accuracy and validity of the genetic variants identified (Price et al. 2006).

Most association studies in natural populations have been focused on candidate immunity-related genes such as the major histocompatibility complex (MHC), Toll-like receptors (TLRs), or cytokines, particularly within single host-parasite systems (compiled by Cheynel et al. 2023). The advancement in next-generation sequencing makes it possible to investigate the host genetic variations across the genome that influence the susceptibility to infections in natural populations beyond the immunity loci. Despite this advancement, genome-wide association studies (GWAS) across spatially heterogeneous populations remain rare in wild animals due to the strong population structure and relatedness among individuals that are commonly found in natural populations confounding association signals (Santure & Garant, 2018). However, relatively few studies have applied GWAS to structured wild populations e.g., Cornetti & Tschirren (2020) used restriction site-associated DNA sequencing (RAD-seq), including both the GWAS and Fst outlier approaches, to identify SNPs associated with parasite-mediated selection imposed by *Borrelia* infections in bank voles across Switzerland (Cornetti & Tschirren, 2020).

Here, we investigate genetic associations between host genome-wide variants with infection by multiple parasites taxa in natural populations of bank voles *Myodes (=Clethrionomys) glareolus* (Schreber, 1780) (*Arvicolinae*, *Rodentia*). Bank vole population structure in Europe have been shaped by repeated colonisation waves from the glacial refugia and ongoing admixture, resulting in higher genetic diversity at mid-latitudes (Marková et al. 2020). Using genome-wide SNP data and a mixed-model association approach that accounts for population structure, we examine genotype-infection associations across ten populations spanning Central Europe. By combining multiple parasite groups within a single host species, our study aims to identify candidate loci involved in susceptibility to infection and to assess the role of parasites in shaping host genomic variation in a spatially structured natural system.

## Materials and Methods

### Data collection and parasite screening

We sampled 182 voles at ten locations (Figure 1a, Table S1) between 2016 and 2023. Although all locations were sampled in late August-September, we found striking differences in vole densities between sites: for instance, in 2023, population density in central Poland (site Chęciny) and in southern Poland (site Beskidy) was below 13 voles per 10^4^ trap-hours; in the same field season with the same trapping effort, it was over 60 in Křižanovská vrchovina (sites Smrk, Vladislav, and Studenec), reaching almost 150 in Březník.

**Figure 1.**
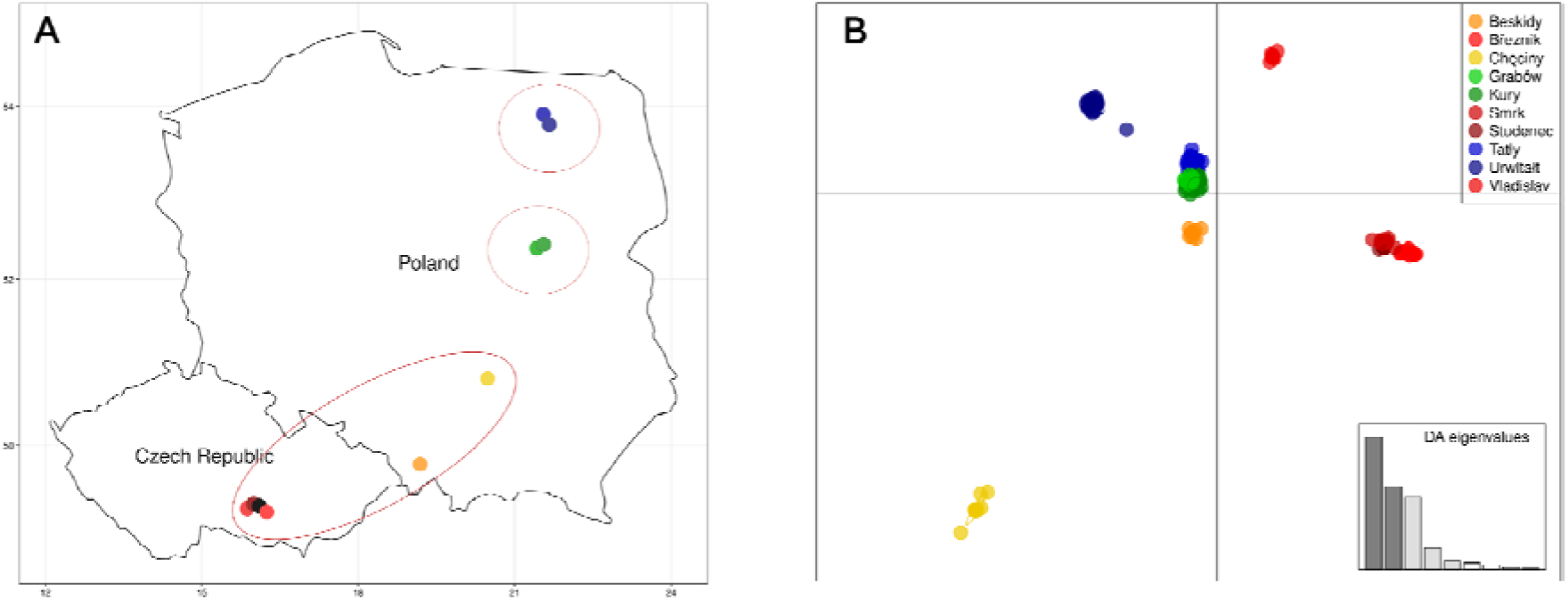
a) Location of the sampling sites. b) Genetic distance between the sampling sites visualized using DAPC (Jombard 2008). PC1 and PC2 are shown.

The collection of material followed the European ethical standard (Directive 2010/63/EU). Voles were euthanized using isoflurane overdose and cervical dislocation, and then weighed and sectioned with the internal organs extracted. They were examined for intestinal parasites, and parasites present were identified morphologically and preserved in 70% ethanol in a pre labelled 1.5ml tube. Spleen was collected for identification of the blood parasites. Material was transported to the laboratory and stored in a -20^0^C.

### Molecular screening of blood pathogens in bank voles

Genomic DNA was extracted from ≤10 mg spleen or liver tissue using the Monarch® Genomic DNA Purification Kit (New England Biolabs), according to the manufacturer’s protocol tissue samples. Each sample was screened for five blood pathogens, namely, *Anaplasma spp*., *Babesia spp*., *Bartonella spp., Rickettsia spp.* and *Toxoplasma spp* using pathogen-specific primers. For details see Table S2. For amplification, we used the OptiTaq polymerase master mix (EurX, Poland); reaction mix included 1.25 U OptiTaq DNA Polymerase, 1 x Reaction Buffer (1.5 mM MgCl_2_), 0.2 mM of each dNTP, 0.3 μM of each primer, and ≤ 100ng of template DNA. For each of the screened pathogens, a previously identified species of each genus was used as a positive control. The PCR products were visualised on 1.5% agarose gels in 1× TAE buffer, run at 120 V for 30 minutes, and stained with SimplySafe (EurX).

### QuaddRADseq library preparation, raw read processing, mapping and SNP calling

Genomic DNA was extracted from ear tissue using the Monarch® Genomic DNA Purification Kit (New England Biolabs), following the manufacturer’s protocol for tissue samples. DNA concentration was measured with the QuantiFluor® ONE dsDNA System on a Quantus™ Fluorometer (Promega). Sample integrity was verified via 1% agarose gel electrophoresis stained with SimplySafe™ (EurX), and only high-molecular-weight DNA was used for library preparation. The adapters used for library preparation included nine quaddRAD_i5n and nine quaddRAD_i7n adapters that were previously used by Martin Cerezo et al. (2022). It supports the restriction enzymes SbfI and MseI and its respective adapter overhangs. For each *M glareolus*, 150 ng of gDNA was simultaneously digested and ligated with adapters in a single step of 40 μL reaction. The reaction mix consisted of 4 μL 10x CutSmart buffer (NEB), 1.5 μL Mse1 (10 UμL^-1^), 0.75 μL Sbf1 (20 U μL^-1^), 4 μL ATP (10mM), 1 μL T4 DNA ligase (400 U μL^-1^), 0.75 μL of each quaddRAD_i5n and quaddRAD_i7n adapter (10 μM) and ddH2O to a final volume of 40 μL. The reaction mixture was incubated for three hours at 30°C in a thermocycler and stopped with 10 μL of 50 mM EDTA. Samples were pooled in sets of 8-9 using dual-index barcodes and double-size selection with the ProNex® Size-Selective Purification System (Promega): a 1:1 (v/v) ratio removed fragments >700 bp, followed by a 0.4:1 (v/v) ratio to remove fragments <250 bp. DNA was eluted in 30 μL of 10 mM Tris (pH 8.5) and quantified using the Quantus™ system. Indexed Illumina adapters were added during a 14-cycle PCR using NEBNext Ultra II Q5 Master Mix. PCR products were purified again using ProNex® (1.5:1 v/v) to remove short fragments (<250 bp), washed, air-dried, and eluted in 20 μL elution buffer. Final DNA concentrations were verified with Quantus™, and fragment distributions assessed on a BioAnalyzer (Agilent Technologies). Libraries were pooled equimolarity, size-selected to 300–600 bp using the Pippin Prep system (Sage Science) at the institute of Nature Conservation, Polish Academy of Sciences, Poland, and sequenced on an Illumina NovaSeq X platform (150 bp paired-end) at Genomed S.A., Warsaw, Poland.

The quality of the raw sequencing reads were assessed and visualized using FastQC v0.12.1 (Andrews, 2010). To remove PCR duplicates, reads were processed using the clone_filter module in Stacks v2.68 (Catchen et al., 2011; Catchen et al., 2013), which identifies duplicates based on four degenerate bases incorporated during quaddRAD library preparation (Tin et al. 2015; Schweyen et al. 2014). Reads were then trimmed using Fastp v0.23.4 to remove low-quality bases (option -q), poly-G tails (option -g), and adapter contamination by specifying expected adapter sequences (Chen et al., 2018; Chen et al., 2023). Demultiplexing was performed using process_radtags in Stacks. During this step, reads with uncalled bases (Ns), low-quality scores (Phred < 20), or that failed Illumina’s chastity filter were discarded. A second FastQC run was performed on the trimmed output to validate filtering effectiveness.

To identify single nucleotide polymorphisms (SNPs), quality-filtered reads were aligned to the *M. glareolus* reference genome using BWA-MEM v0.7.8 (Li, 2013) with default parameters. The resulting SAM files were converted to sorted and indexed BAM files using SAMtools v0.1.18 (Li et al. 2009). SNP calling was performed using the ref_map.pl pipeline in Stacks v2.68, which constructs a catalogue of loci from the aligned reads. The gstacks module was used to generate genotype likelihoods and assess sequencing depth across samples. Samples with an average sequencing depth below 3× were excluded from further analysis. Variant calling was completed by running the populations module in Stacks without applying additional filtering, producing an initial VCF file for downstream population genomic analyses.

### Association analysis

The resulting VCF was filtered in PLINK v 1.9 (Purcell et al. 2007; Chang et al. 2015), removing variants with missing rates of over 10%, variants not in Hardy-Weinberg equilibrium with a p<10⁻⁵ threshold, and variants with minor allele frequencies below 5%. We also filtered out variants in linkage disequilibrium using the --indep-pairwise option with a 150bp window size, a 20 bp step, and a 0.7 threshold r² value. The filtering resulted in 30206 variants in 182 individuals that were used in the GWAS analysis.

The associations between SNP variants and the presence/absence of pathogens were analysed using GEMMA, genome-wide efficient mixed-model analysis for association studies (Zhou & Stephens 2012). Mixed linear models have been shown to effectively account for relatedness among samples and in controlling for population stratification and other confounding factors (Kang et al. 2010, Price et al. 2010). GEMMA controls for population structure and family relatedness using the genetic relationship matrix (GRM), computed using the -gk 1 option, which generates a centred kinship matrix based on genome-wide SNP genotypes. In each model, as a cofactor we used host body mass which has been shown in previous studies to affect the parasite load in bank voles (Kloch et al. 2018). In the GWAS analysis, we included parasites with a total prevalence over 10%. Additionally, for each pathogen, we run separate models with the sampling site as an additional covariate.

The results were controlled for multiple comparisons using the conservative Bonferroni correction. For each model, we calculated the genomic inflation factors (λ) to assess the effectiveness of the structure correction (Devlin & Roeder, 1999; Price et al. 2010). Models with λ close to 1 indicated no inflation, suggesting that population structure and relatedness were adequately accounted for. In contrast, high λ values indicated inflation due to confounding factors such as family relatedness, and in some cases suggested that the trait of interest may be polygenic (Yang et al., 2011).

To describe the genetic structure, we used DAPC (discriminant analysis of the principal components, Jombart et al. 2010) implemented in the R package adegenet (Jombart, 2008). The method allows for the identification of clusters of the most genetically similar individuals, maximising variance between groups and minimising variance within groups. The DAPC was also used for summarising genetic differences between sampling locations.

### Gene and gene ontology annotation in associated SNPs

SNPs identified from the RADseq dataset were functionally annotated using SnpEff v5.2 (Cingolani et al. 2012). The *M. glareolus* reference genome (Marková et al. 2023), together with its corresponding GFF annotation file, was used to build a custom SnpEff database. with respect to the annotation in the reference genome. The VCF containing the filtered SNPs was processed with the custom database to extract the effects of each variant. For the SNPs associated with infection risk, the gene name, gene function, gene Ontology (GO) annotations, and associated GO terms of the corresponding gene were retrieved from the UniProt Knowledgebase (UniProtKB; The UniProt Consortium, 2025 https://www.uniprot.org/). Also, we used 10 kb sequences around the significant SNPs to search the NCBI nucleotide database to identify homologous genomic regions and further confirm the functions of the genes on the Uniprot Knowledgebase. There were some differences among methods reflecting differences in annotation depth and homology detection, but the majority of the genes overlapped (see Table S3).

## Results

### Spatial heterogeneity

The overall parasite prevalence ranged between 13% (*Aonchotheca annulosa*) to 38.4% (*Heligmosomum mixtum*), and the differences between sites were significant for all parasites except for the least common *A. annulosa;* for example, the *H. mixtum* infected >50% of examined voles in central and north Poland (sites Kury, Grabów, Urwiłtat, Tałty) but was absent from several sites in the Czech Republic (Březník, Studenec, and Vladislav) (Table S1). These differences could not be explained by the differences in host density for any of the parasites, as all correlations between prevalence and density were weak and not significant (Table S4).

Discriminant analysis of the principal components (DAPC) revealed that studied populations are highly structured. Although the data comprised 10 sampling sites from five regions, the most likely number of clusters, as indicated by the lowest BIC score, was 3. Cluster 1 grouped sites in southern Poland and the Czech Republic (Figure 2), although the Sudetes mountain range separates them, and they differ considerably in the parasite loads. Cluster 2 consisted of two sites in northern Poland, and cluster 3 included two sites from central Poland. The loci that contributed most to that division were Scaffold_4289;HRSCAF=4559, ID 300295:17, followed by Scaffold_4285;HRSCAF=4508, ID 60876:28 and Scaffold_3734;HRSCAF=3925, ID 612472:80.

**Figure 2.**
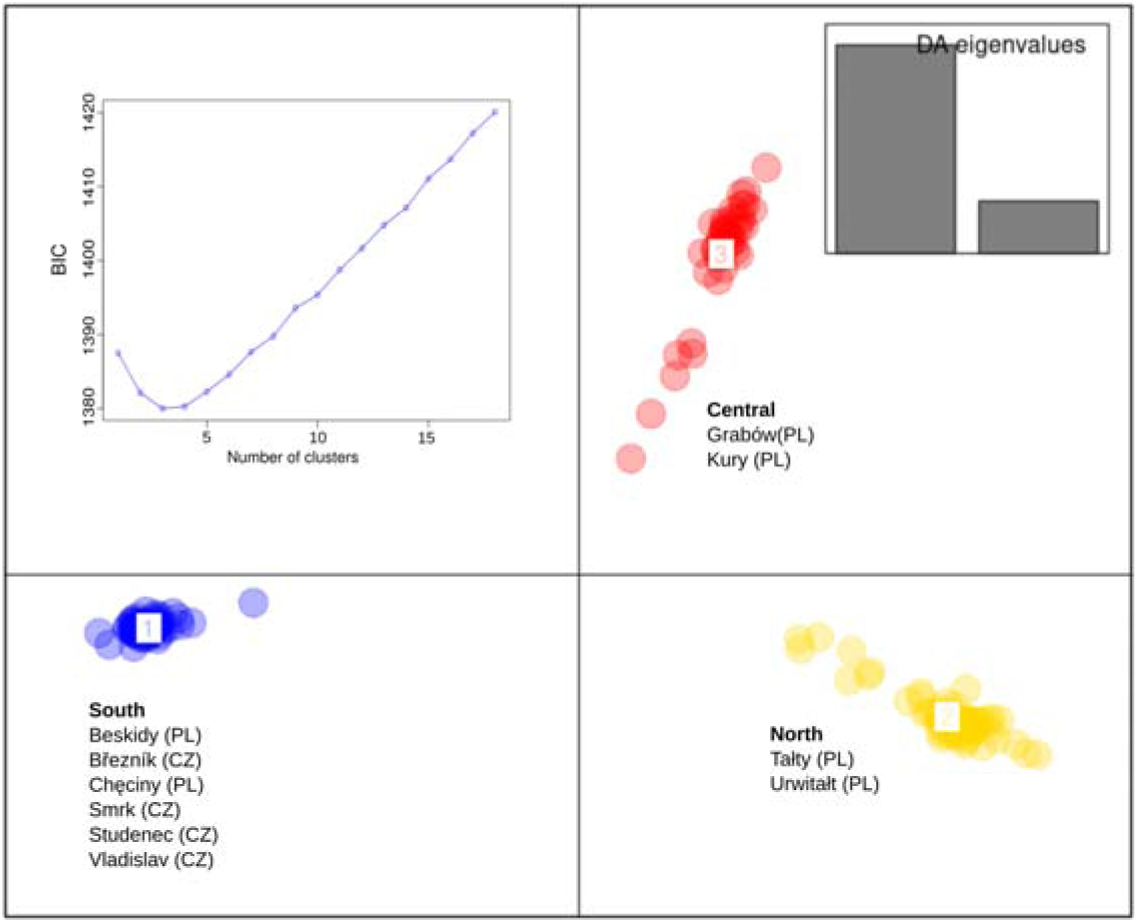
Genetic structure of the samples with no prior assumption on their membership analysed using DAPC. The most likely number of clusters was 3, as indicated by the BIC plot. By each cluster represented in different colour we present a list of sampling sites grouped in this cluster.

When we applied DAPC to actual sampling sites (Figure 1b), the site Chęciny in central Poland appeared to be the most distant from the rest. There was a visible trend along PC2 from north - sites Tałty and Urwitałt, marked in Figure 1b in yellow - to the south (sites Březník, Vladislav, Studenec, and Smrk marked in blue). The loci that contributed most to that division were Scaffold_971;HRSCAF=1034, ID 398766:82 and Scaffold_2945;HRSCAF=3096, ID 548523:23 followed by Scaffold_971;HRSCAF=1034, ID 383070:50

### Loci associated with susceptibility to helminth and blood infections

We identified twelve loci significantly associated with susceptibility to helminth and blood-borne infections (Table 1, models for all the loci and all the parasites are presented in Table S5 and Table S6). Out of the ten parasites and pathogens tested, we found SNP exceeding the genome-wide significance threshold based on likelihood-ratio tests with significance level adjusted for multiple comparisons in six taxa. The genomic inflation value (λ) for the models for each taxon ranged between 0.80 and 1.06 that indicates no inflation, (Table S7) i.e. that the models adequately accounted for population structure and relatedness between analysed individuals.

**Table 1:** Loci significantly associated with the susceptibility to helminth and blood infections detected using Gemma. OR - odds ratio, β - parameter estimate, p-val - p-value of the association based on likelihood ratio test. Results for all SNPs and all parasites are given in Table S5.

| parasite | chromosome | position | SNP ID | minor allele | major allele | minor allele freq | OR | $\beta$ | p-val (LR) | $\beta$ (sites as cofactor) | p-val (LR) (sites as cofactor) |
| --- | --- | --- | --- | --- | --- | --- | --- | --- | --- | --- | --- |
| *A. tianjinensis* | Scaffold_4289;HRSCAF=4559 | 80398611 | 342121:13 | T | C | 0.057 | 4.727 | -0.376 | $9.8 \times 10^{-7}$ | -0.371 | $1.8 \times 10^{-6}$ |
| Cestodes | Scaffold_4288;HRSCAF=4551 | 25076629 | 856263:58 | T | C | 0.061 | 0.149 | 0.426 | $2.81 \times 10^{-6}$ | 0.424 | $2.90 \times 10^{-6}$ |
| *H. mixtum* | Scaffold_4285;HRSCAF=4508 | 124979183 | 104360:101 | A | G | 0.06 | 4.496 | 0.887 | $1.03 \times 10^{-8}$ | 0.867 | $5.18 \times 10^{-9}$ |
| *H. mixtum* | Scaffold_4270;HRSCAF=4487 | 23900832 | 1757333:20 | T | C | 0.052 | 0.368 | 0.881 | $5.41 \times 10^{-8}$ | 0.978 | $5.06 \times 10^{-10}$ |
| *H. mixtum* | Scaffold_4300;HRSCAF=4617 | 2903852 | 1298450:110 | A | G | 0.05 | 1.739 | 0.892 | $6.63 \times 10^{-8}$ | 0.924 | $1.43 \times 10^{-8}$ |
| *H. mixtum* | Scaffold_4295;HRSCAF=4597 | 53142752 | 1106417:192 | T | C | 0.066 | 0.36 | 0.745 | $1.34 \times 10^{-7}$ | 0.718 | $2.29 \times 10^{-7}$ |
| *H. mixtum* | Scaffold_971;HRSCAF=1034 | 87023895 | 448318:52 | A | G | 0.05 | 2.222 | 0.787 | $6.67 \times 10^{-7}$ | 0.884 | $1.06 \times 10^{-8}$ |
| *H. mixtum* | Scaffold_4304;HRSCAF=4627 | 29562250 | 1824002:107 | A | G | 0.058 | 1.422 | 0.812 | $7.7 \times 10^{-7}$ | 0.792 | $1.19 \times 10^{-6}$ |
| *H. mixtum* | Scaffold_3734;HRSCAF=3925 | 36525066 | 604510:8 | T | C | 0.072 | 1 | 0.696 | $7.83 \times 10^{-6}$ | 0.717 | $3.45 \times 10^{-7}$ |
| *M. muris* | Scaffold_4295;HRSCAF=4597 | 44438406 | 1098592:169 | T | C | 0.05 | 0.098 | 0.436 | $5.68 \times 10^{-7}$ | 0.4073 | $2.55 \times 10^{-6}$ |
| *Rickteksia* ssp. | Scaffold_1703;HRSCAF=1800 | 2181349 | 1678608:183 | A | G | 0.054 | 0.238 | 0.528 | $1.7 \times 10^{-6}$ | 0.555 | $1.7 \times 10^{-6}$ |
| *Toxoplasma* ssp | Scaffold_3734;HRSCAF=3925 | 108109541 | 664806:134 | T | C | 0.456 | 0.408 | 0.271 | $4.5 \times 10^{-7}$ | 0.2736 | $3.18 \times 10^{-6}$ |

Out of the twelve SNPs identified, ten are significantly associated with helminth infections, one with the presence of bacteria *Rickettsia*, and one with the presence of the protozoan *Toxoplasma*. Interestingly, seven of them are significantly associated with *H. mixtum* infections. Individuals with the minor allele in most of the significant SNPs were less likely to be infected, and the strongest effects were observed across the SNPs associated with the presence of infection with *H. mixtum* (0.696 - 0.892), with SNPs 1298450:110 (OR= 1.739, β=0.892) with the strongest effect. On the contrary, the SNP on the Scaffold_4289;HRSCAF=4559 (position 80398611) was associated with *A. tianjinensis* and individuals with the minor allele in the locus 342121:13 were 4.72 times more likely to be infected with the *A. tianjinensis*. Minor allele frequencies were generally low (∼5-6%), except for the locus associated with *Toxoplasma sp.*, which involved a common allele (MAF = 0.456) (Table 1).

The same set of significant SNPs were associated when sites were included as a cofactor, with only slight differences in the values of the effect sizes indicating that these associations are robust to spatial structure (Table 1, Table S6). The only exception was *H. mixtum* - in this case the analysis revealed significant associations between 38 SNPs and susceptibility to infection. Among them were the 6 loci reported in models without sites as a cofactor.

### Candidate genes for susceptibility to infections

SnpEff annotations classified the identified twelve SNPs as MODIFIER variants as they are intergenic, intronic, or upstream or downstream gene variants, with none predicted to alter coding sequences (Table S3 and S8). Two loci were within 10 kb up- or downstream of protein-coding genes, including *Pi4k2a*, which was associated with cestode infection, and *Rabep1* and *Elmod3*, which were associated with *H. mixtum* infection. Four other SNPs are between 17kb and 43kb of Nuak2, Galnt14, Mapkapk2, Pol genes. And the other SNPs were found above 100kb of Ntf3, Kcna5, Ikzf5, Cdkal1, Nuak1, Lclat1, Fastkd2, Pcdh9, Tnni3k, Eef1e1, homology genes in *Mus musculus* (Table 2). They are all involved in diverse molecular functions and biological processes mostly including catalytic activity, metal ion binding, cellular component organisation, signalling and response to stimulus (Table S5 and S6). They are highly conserved across animal taxa and represent diverse functional pathways, including phosphoinositide and lipid signalling, vesicle trafficking, and cellular stress responses.

**Table 2.** Genes in proximity to significant intragenic loci detected as significantly associated with infections in Gemma.

| chromosome | position | SNP ID | variant type | parasite | gene | distance of gene to variants (bp) |
| --- | --- | --- | --- | --- | --- | --- |
| Scaffold_3734;HRSCAF=3925 | 108109541 | 664806:134:+ | intergenic | *Toxoplasma* | Galnt14<br>Lclat1 | 43430 to 40519,<br>-216216 to -216674 |
| Scaffold_1703;HRSCAF=1800 | 2181349 | 1678608:183:+ | intergenic | *Rickettsia* | Nuak1<br>Nuak2 | 147239 to 82420<br>-26664 to -27107 |
| Scaffold_4288;HRSCAF=4551 | 25076629 | 856263:58:- | downstream gene variant | Cestodes | Pi4k2a | -2705 to -3198 |
| Scaffold_4289;HRSCAF=4559 | 80398611 | 342121:13:- | intergenic | *A. tianjinensis* | Pik3r1 | 137103 to 136763 |
| Scaffold_4295;HRSCAF=4597 | 44438406 | 1098592:169:+ | intergenic | *M. muris* | Ikzf5<br>Cdkal1 | -539504 to -692097 |
| Scaffold_4285;HRSCAF=4508 | 124979183 | 104360:101:- | intergenic | *H. mixtum* | Fastkd2 | -404631 to -407298 |
| Scaffold_4270;HRSCAF=4487 | 23900832 | 1757333:20:+ | intergenic | *H. mixtum* | Pcdh9 | -2034199 to -2034587 |
| Scaffold_4300;HRSCAF=4617 | 2903852 | 1298450:110:- | intergenic | *H. mixtum* | Tnni3k | -1064707 to -1313827 |
| Scaffold_4295;HRSCAF=4597 | 53142752 | 1106417:192:+ | intergenic | *H. mixtum* | Eef1e1 | -335618 to -335938 |
| Scaffold_971;HRSCAF=1034 | 87023895 | 448318:52:+ | intergenic | *H. mixtum* | Mapkapk2 | 33175 to 28078 |
| Scaffold_4304;HRSCAF=4627 | 29562250 | 1824002:107:- | intergenic | *H. mixtum* | Pol | -17859 to -18162 |
| Scaffold_3734;HRSCAF=3925 | 36525066 | 604510:8:+ | upstream gene variant<br>(transcript/ protein_coding) | *H. mixtum* | Rabep1<br>Elmod3 | 3274 to 2902 |

## Discussion

Understanding host susceptibility to pathogens is essential for explaining how evolutionary pressures maintain diversity in resistance and susceptibility traits within and among natural populations, as well as for understanding the emergence and re-emergence of infectious diseases (Vicente-Santos et al., 2023; Höckerstedt et al., 2022). Numerous studies confirmed the associations between variation at single loci, often coding for the components of the immune system, and susceptibility to infections in free-living species. In bank voles, susceptibility to nematodes was shown to be associated with SNP in cytokines and MHC genes (Kloch et al. 2010, 2023). Similarly, associations between susceptibility to bacterial infections and variation in toll-like receptor genes have been reported (Tschirren et al. 2013; Kloch et al. 2018). However, contrasting evidence was found by Tarnowska et al. (2020), with no association between genetic variation in toll-like receptor genes and *Borrelia afzelii* infection prevalence in north-eastern Poland. In this study, we performed a genome-wide association study (GWAS) to identify single nucleotide polymorphisms (SNPs) associated with susceptibility to both helminth and blood pathogens in a highly structured wild populations of the bank vole (*M. glareolus*).

### Loci associated with susceptibility to infections in the bank vole

We identified twelve candidate host loci associated with host genetic susceptibility to helminth and blood-borne pathogen infections in wild rodents at inter-population level. Of the identified variants, ten were located in intergenic regions. The remaining two variants, were located near protein-coding genes. One SNPs associated with cestodes is downstream of *Pi4k2a* and the other associated with *H. mixtum* is upstream of *Rabep1* and downstream of *Elmod3*. The genes associated with the identified variants are highly conserved across animal taxa and represent diverse functional pathways, including phosphoinositide and lipid signalling, vesicle trafficking, transcriptional regulation, and cellular stress responses. Together, these functions indicate that host susceptibility to helminths and blood pathogens is influenced not only by known immune responses but also by regulatory, metabolic, and membrane-associated processes which could affect host tolerance to infection and parasite persistence.

*H. mixtum* infection was associated with the highest number of significant loci. Among these are two variants located within 29kb of *Mapkapk2* and 3kb of *Rabep1* genes. These genes are implicated in functions relevant to host-parasite interactions, including immune signaling, vesicle trafficking, and response to cellular stress. *MAPKAPK2 encodes* MAP kinase-activated protein kinase 2, a serine/threonine-protein kinase activated by MAP kinase p38-alpha/MAPK14 (Kotlyarov, et al. 1999; Stokoe et al. 1993). MAPKAPK2 is involved in multiple cellular processes, including cytokine production, endocytosis, and Toll-like receptor (TLR) signaling in dendritic cells (Menon, et al. 2013: Zaru et al. 2007). Inhibition of p38 MAPK signaling impairs host defense mechanisms, including Th1-mediated immunity against intracellular parasites such as *Leishmania* (Yang et al. 2010), and reduces autophagy-mediated bacterial clearance, resulting in increased intracellular *Salmonella* loads in mouse embryonic fibroblasts (Suwandi et al. 2023). The proximity of a SNP associated with increase with susceptibility to *H. mixtum* infections to *Mapkapk2* suggests that variation in this genomic region may be linked to pathways involved in immune activation or cellular stress responses during infection. However, the associated variant lies outside the coding sequence and may represent a regulatory polymorphism. Similarly, an upstream gene variant near *Rabep1* was associated with susceptibility to *H. mixtum* infection. RABEP1 (RAB GTPase Binding Effector Protein 1) is a member of the Rab family of small GTPases, which regulates intracellular membrane trafficking in immune cells during inflammation (Homma et al. 2021; Prashar et al. 2017). Rab GTPases control phagosome maturation, endocytosis and exocytosis and pathogens such as *Legionella*, *Salmonella* exploit these pathways to survive intracellularly (Flannagan et al. 2012; Prashar et al. 2017; Spanò et al. 2016). These functions make the region near *Rabep1* also a biologically plausible candidate involved in host-parasite interactions.

The susceptibility to presence of Cestodes was linked to a downstream gene variant within 3kb of Pi4k2a gene. The gene encodes phosphatidylinositol 4-kinase 2-alpha, an enzyme that catalyzes the phosphorylation of phosphatidylinositol (PI) to phosphatidylinositol 4-phosphate (PI4P) upstream of PI3K/AKT signalling (Guo et al. 2003; Chu et al. 2010). PI4P is a lipid signalling molecule involved in Golgi organization, endocytosis, protein sorting, and membrane trafficking (Barylko et al. 2001; Minogue et al. 2001; Minogue et al. 2010; Mössinger et al. 2012). The gene is evolutionarily conserved among eukaryotes, in *Rattus norvegicus*, it has been shown to regulate Ca²⁺-dependent exocytosis in mast cells (Ishihara et al. 2006). Notably, several pathogens including parasites, viruses and bacteria are able to replicate and survive intracellularly through the mechanisms they have developed to exploit the host PI4Ks (Mendes et al. 2025). Although the associated SNP lies outside the coding region and may represent a regulatory variant, its close proximity to Pi4k2a makes this region a plausible candidate for involvement in host-parasite interactions.

Several other SNPs were located more distantly from annotated genes (≥17 kb up to >500 kb), e.g. a SNP associated with susceptibility to *A. tianjinensis* was located in the intergenic region approximately 228 kb upstream of Ntf3 (Neurotrophin-3) and 201 kb downstream of Kcna5 (Potassium Voltage-Gated Channel Subfamily A Member 5). Ntf3 encodes neurotrophin-3, a growth factor that signals through NTRK3 receptors, activating PI3K/AKT signalling and PLCγ pathway involved in cell survival and differentiation, stress responses, and tissue homeostasis (Marsh & Palfrey 1996; Liot et al. 2004). It has been suggested that regulatory variants near Ntf3 could affect host tissue homeostasis during an infection (Liot et al. 2004). Another locus located in close proximity to Ikzf5 and Cdkal1 was significantly associated with infection by *M. muris*. *Ikzf5* (Ikaros family zinc-finger protein 5, also known as Pegasus) is a member of the Ikaros family of zinc-finger transcription factors and is expressed in lymphocytes, where it has been implicated in the regulation of lymphoid development (OMIM, 2002). Although *IKZF5* has been less extensively studied than other family members (*IKZF1-3*), the Ikaros family as a whole plays a central role in immune regulation, particularly through control of cytokine signalling pathways and CD4⁺ T-helper cell differentiation (Powell et al. 2019). *Cdkal1*, located near the same variant, is a member of the methylthiotransferase family and encodes CDK5 (Ching et al. 2002). Genetic variation in *Cdkal1* has been associated with changes in insulin secretion and glucose metabolism and its deficiency causes pancreatic islet hypertrophy in mice (Wei et al. 2011). Chang et al. (2014) proposed that together with *GIP, Cdkal1* may be under environmental selection and may indirectly affect adaptive immunity.

Among the blood-borne pathogens, only two loci were significant. A SNP associated with *Rickettsia* spp. infection was located approximately 83 kb upstream of *Nuak1* and 27 kb downstream of *Nuak2*, both members of the SNF1/AMPK-related kinase family. They participate in cellular stress responses and regulate processes such as cell adhesion, proliferation, and survival (Humbert et al. 2010; Suzuki et al. 2004; Zagórska et al. 2010). NUAK1 has been shown to directly phosphorylate and modulate the activity of p53/TP53, thereby influencing cell survival under metabolic stress through regulation of DNA damage responses and apoptosis (Hou et al. 2011). The associated variant identified here lies in a non-coding region and may reflect linkage to regulatory elements rather than direct functional variation. The SNP associated with *Toxoplasma* spp. infection mapped within *CAPN13* and was located 43 kb from *GALNT14* and 216 kb from *LCLAT1*. *GALNT14* encodes a glycosyltransferase involved in mucin-type O-glycosylation (Wang et al. 2003) and the loss of GALNT14 function connects to IgA nephropathy due to excess IgA production, impaired B lymphocyte homing, and altered mucosal immunity (Pell & Menon, 2025; Prakash et al. 2025). While *LCLAT1* encodes lysocardiolipin acyltransferase 1, involved in phosphatidylinositol acyl-chain remodeling (Zhao et al. 2009; Bone et al. 2017). Bone et al. (2017) demonstrated that disruption of Lclat1 leads to defects in endocytosis and endocytic trafficking. It is also required for effective PI3K-Akt signaling downstream of receptor tyrosine kinases (Chan et al. 2024). CAPN13 is a member of the calpain large subunit family of calcium-dependent cysteine proteases activated by intracellular Ca²⁺ signaling (Dear & Boehm, 2001) and are implicated in intracellular signaling and cytoskeletal dynamics.

Several annotated genes located more distantly away from the associated SNPs (≥17 kb up to >500 kb) are involved in processes such as cellular stress responses and tissue homeostasis, cytokine signalling, CD4⁺ T-helper cell differentiation, endocytosis and endocytic trafficking. These pathways are relevant to immune functions. However, we can not confirm the link between the genetic variants and these genes. It is possible that some of these variants influence host resistance and susceptibility through long-range regulatory mechanisms, such as having effect on gene expression or chromatin interactions, as long-range regulatory relationships have been documented in vertebrate genomes (Chandra et al. 2021; Mercurio et al. 2023). The confirmation of this relationship is beyond the scope of this study. Future studies integrating functional genomics approaches will be necessary to determine whether these distal variants have regulatory effects relevant to host-parasite interactions.

### Effect of population structure on the GWAS outcome

Controlling for population structure is crucial in studies from free-living species that usually tend to be structured due to environmental heterogeneity and complex demography. Failure to account for population structure could cause spurious associations, resulting in false positives that can bias the GWAS results (Bouaziz et al. 2011; Liu et al. 2023). However, Gloss et al. (2023) revealed that while correcting for population structure is effective at reducing false positives, these corrections can reduce the power to detect true causal variants that also vary geographically. Similarly, Wang et al. (2021) investigated the impact of not controlling for population structure in an inbred mouse strain with known causative traits and found that population structure correction could result in rejection of association signals that were generated by these known causative alleles.

Previous work in free living *M. glareolus* did not detect significant SNP associations with *Borrelia afzelii* infection after correcting for multiple comparisons (Cornetti & Tschirren 2020), even though the authors controlled for the population structure. They however identified several candidate loci using the Fst outlier method. Zhao et al. (2007) showed that the algorithm used for GWAS testing may fundamentally alter the results, and in most cases only mixed-models can adequately account for the population structure. Our study confirms this finding. We used univariate linear mixed models (LMM) implemented in GEMMA, and only this method resulted in genomic inflation value (λ) within acceptable limits. When different method was applied i.e. when population structure was included in the models as a cofactor rather than fixed effect using R package SNPassoc (González et al. 2007), the λ was close to 2, indicating that the models failed to account correctly for the variation in data associated with kinship between individuals and population structure (results not shown).

In the current work we did not detect any of the associations between variation at immunity loci and susceptibility to infections reported in our previous works (Kloch et al. 2010, Kloch et al. 2018, Kloch et al. 2023). These previous studies were restricted to a single geographic region, namely 3 populations located in NE Poland, two of which (Urwitałt and Tałty) were also included in the current study. They showed that even in such a small scale, within ∼20km distance, the associations between alleles and susceptibility may have contrasting effects between locations: an allele that in one location was associated with higher resistance, in another was linked to higher susceptibility (Kloch et al. 2010). In the present work, the geographic scale was much larger, comprising 10 locations spread longitudinally across ∼750km. Since the parasite communities tend to differ between locations (see Table S1), they constitute a spatially heterogeneous selective force. As a consequence, the local parasite-driven co-evolution resulting in spatially contrasting effects of alleles might have cancelled out and remained invisible for GWAS analysis. Thus, the effects reported here are likely linked to general mechanisms of resistance rather than results of local host-parasite arms race. This emphasizes that not only the way of controlling spatial structure, but also the spatial scale, may affect the outcome of the GWAS analysis. The associations detected in the current work likely represent a general mechanism of pathogen resistance rather than results of parasite-driven local selection.

In our genome-wide association analyses of five blood pathogens and five helminth taxa, only two pathogens and three helminth taxa was associated with single genome-wide significant locus each, one helminth (*H. mixtum*) was associated with seven significant loci. The five remaining taxa had no significant loci above the Bonferroni correction. These are not unexpected in GWAS of natural host-parasite systems, where limited sample sizes and complex genetic architectures reduce statistical power to detect individual loci with small effects (e.g., Santure and Grant, 2018; Spencer, et al. 2009; Visscher et al. 2017). In addition, stringent multiple-testing corrections required in GWAS to control false positives can further diminish power (Manolio et al., 2009; Bush and Moore, 2012).

In host-parasite systems, effect sizes are often small and distributed across many loci, consistent with a highly polygenic interaction landscape that is difficult to resolve with limited sample sizes (Santure and Garant 2018; Zhang et al., 2018). This pattern was observed in *H. mixtum* for which we identified seven loci distributed across different scaffolds to be significantly associated with host genetic variation. However, RAD-seq has limitations for GWAS due to its sparse and uneven coverage of the genome, as it only captures loci near restriction sites (e.g., Lowry et al., 2017). Future studies could improve resolution and power by using low-coverage whole-genome sequencing, which allows more comprehensive variant discovery and denser marker coverage (e.g., Chat et al., 2022; Lou et al. 2021).

To improve discovery in future studies of taxa that did not yield any significant SNPs, we recommend complementing GWAS with secondary analyses that are more sensitive to polygenic signals and selection signatures. For example, combining association tests with population differentiation (Fst or outlier) scans (Cornetti and Tschirren, 2020), incorporating gene expression after selecting highest differentiated SNPs (Queirós et al. 2018), or considering reporting of exploratory unadjusted p-values and independent replication where feasible (Elbers et al. 2018; Pérez-Umphrey et al. 2023). Combining complementary approaches may help reveal host genetic factors that are otherwise obscured by limited power in genotype-phenotype GWAS.

To our knowledge, this is the first GWAS to simultaneously assess susceptibility to multiple parasite types (helminths, protozoa, and bacteria) in a free-living host species across spatially heterogeneous populations. We identified several novel candidate genes associated with susceptibility or resistance to infections. The identified loci have overlapping pathways, including stress signaling (e.g., MAPKAPK2, NUAK1/2), vesicle or membrane trafficking (e.g., RABEP1, LCLAT1, PI4K2A), and immune regulation (e.g., IKZF5, CAPN13, NTF3). Most of the variants lie in regulatory regions, suggesting that variation in gene expression, other than changes in protein coding sequences, may also be important in differences in susceptibility of host to infections. Overall, these findings demonstrate the importance of accounting for both well-known and less obvious genes in studies of host-parasite interactions and the evolution of disease resistance.

## Supporting information

Supplementary_TablesS1-S4andS7-S8

## Acknowledgements

We appreciate the support of Joelle Gouy de Bellocq and Alena Fornůsková from Czech Academy of Sciences, Institute of Vertebrate Biology for their help during data collection in the Czech Republic, Jarosław Bryk for access to the Orion computer cluster during AEO research visit at University of Huddersfield, Anna Karnkowska from University of Warsaw for access to the server at the Institute of Evolutionary Biology, Aleksandra Biedrzycka and Joanna Kołodziejczyk, for help with the use of the Pippin Prep machine at the Institute of Nature Conservation, Polish Academy of Sciences.

## Data Accessibility and Benefit-Sharing section

### Data Accessibility Statement

The ddRAD sequencing data generated and analyzed in this study, together with additional related samples from the broader project, are available through the National Center for Biotechnology Information’s Sequence Read Archive under BioProject accession PRJNA1465137. The filtered SNP dataset, codes and related metadata supporting the findings of this study have been made publicly available in Zenodo (DOI 10.5281/zenodo.20271351). The metadata include sampling georeferences in decimal degrees, collection dates (day/month/year), infection status of individual bank voles to the different parasites screened and each BioSample accession linking both the deposited genetic data and deposited metadata.

### Benefit-Sharing Statement

We consulted local scientists during sampling collection, their the contributions have been described in acknowledgements. All data have been shared with the broader public via appropriate biological databases as described above.

### Author’s contributions

A.E.O., Methodology, Data collection, Investigation, Data analysis, Funding Acquisition, Writing – Original Draft, Manuscript review and editing; J.B., Methodology, Data analysis, Writing – Original Draft Manuscript review and editing; V.I.A., Data collection, Investigation, Manuscript review; A.D., Data collection, Investigation; A.R., Data collection; A.K., Conceptualisation, Methodology, Data collection, Data analysis, Funding Acquisition, Writing, Manuscript review and editing, Supervision. All authors gave final approval for publication.

### Financial Support

This study was funded by grant 2020/39/O/NZ8/01596 from the Polish National Science Foundation (NCN) to A.K., grant BPN/PRE/2022/1/00025/U/00001 from the Polish National Agency for Academic Exchange and BOB-IDUB-622-479/2023 micro-grant under Action IV.4.1 “A complex programme of support for UW PhD students” implemented in the program “Excellence Initiative – Research University” to A.E.O.

### Ethical standards

All procedures involving animals were reviewed and approved by the National Ethical Committee for Research on Animals under Resolution No. 12/2022. Fieldwork was conducted in state-owned forests, and no specific permits or licences were required for sampling.

